# Temporal Focusing for Enhanced Background Rejection in AOD-Based Two-Photon Serial Holography

**DOI:** 10.64898/2026.03.07.710267

**Authors:** Joséphine Morizet, Walther Akemann, Benjamin Mathieu, Jean-François Léger, Laurent Bourdieu

## Abstract

The ability to record 3D neuronal activity with cellular resolution, high signal-to-noise ratio (SNR) and millisecond temporal resolution is a major challenge in neuroscience. One powerful method is random-access two-photon microscopy based on acousto-optic deflectors (AODs), which uses a holographically-shaped point spread function (PSF) scanned in 3D to maximize the sampling rate and SNR. However, this approach suffers from greater background contamination due to the holographically shaped PSF than standard two-photon microscopy with diffraction-limited PSF. To overcome this limitation, we implemented a new version of an AOD scanning system, which integrates temporal focusing. The complex spatiotemporal distortions encountered in this configuration, including a significant group delay dispersion associated with the pulse front tilt generated by the AOD, were compensated for by introducing an acousto-optic modulator before the AOD. We designed extended patterns by combining temporal focusing on one direction and holographic wavefront shaping in the perpendicular axis. Taking advantage of the AOD’s ability to shape the wavefront at the same speed as the scan, we were able to accurately superimpose the spatial and temporal foci over the entire field of view. Finally, we generated complex, extended two-photon excitation patterns by combining temporal focusing in one direction and holographic multiplexing in the perpendicular direction. These patterns provide significantly improved background rejection compared to 2D holographic patterns, thus offering promising prospects for in vivo recordings of neuronal activity in dense samples with improved SNR.

## 1. Introduction

3D optical recording of neuronal activity with cellular resolution within deep tissue and at high spatiotemporal resolution remains a major challenge in neuroscience [1–3]. Non-linear two-photon fluorescence microscopy (TPM) is the method of choice for addressing this question, as it allows images with sub-cellular resolution to be obtained deep within scattering tissues and with excellent optical sectioning. Its resolution and optical sectioning also make it highly sensitive to fluorescence modulation generated by activity indicators expressed in small neuronal compartments (cytoplasm, plasma membrane). TPM and genetically encoded calcium indicators (GECIs) have thus allowed the in vivo measurement of neuronal activity in genetically identified cells in head-fixed or freely moving rodents, by recording calcium transients evoked by action potentials [4]. Recently, advances in genetically encoded voltage indicators (GEVIs) have provided direct access in TPM to actual voltage dynamics, including subthreshold and spike activity [5].

Yet, two-photon recordings of the fast fluorescence signals from these indicators with high signal-to-noise ratio (SNR) and across large neuronal populations remain challenging. The problem is particularly difficult if the neurons of interest are positioned in 3D, e.g. at different depths within a cortical column. A promising solution is the use of random-access microscopy, which allows maximizing sampling rate and SNR by addressing the laser beam only to the cells of interest (see Figure 1a). Random-access is currently most successfully implemented using acousto-optic deflectors (AODs), which enable fast, millisecond range, 3D targeting of cells [6–10]. The required tilt and defocus of the laser wavefront to achieve 3D random access is obtained by offsetting and linearly chirping the carrier frequency of the acoustic wave in the AODs [6]. One implementation of AOD-based 3D random access is 3D Custom Access Serial Holography (3D-CASH) [11]. This approach relies on the synchronization of AODs with the repetition rate of the laser [10], allowing thus the microscope Point Spread Function (PSF) to be shaped by digital holography with the AODs, as well as scanned in 3D (see Figure 1a,b). The holograms (see Figure 1c) are implemented thanks to the use in the AODs of ultrasonic waves modulated in amplitude and frequency, which adds to the AODs the ability to be used as spatial light modulators. [11, 12]. The spatial shaping of the PSF has two aims : to maximize the photon flux from the recorded cells and to mitigate motion artifacts observed in vivo [11]. 3D-CASH microscopy has achieved volumetric recordings of neuronal activity with millisecond resolution in head-fixed behaving mice (see Figure 1b), of typically 100 cells at 400Hz using GECIs [11] or 10 cells at 4kHz using GEVIs (Akemann, in preparation).

**Fig. 1.**
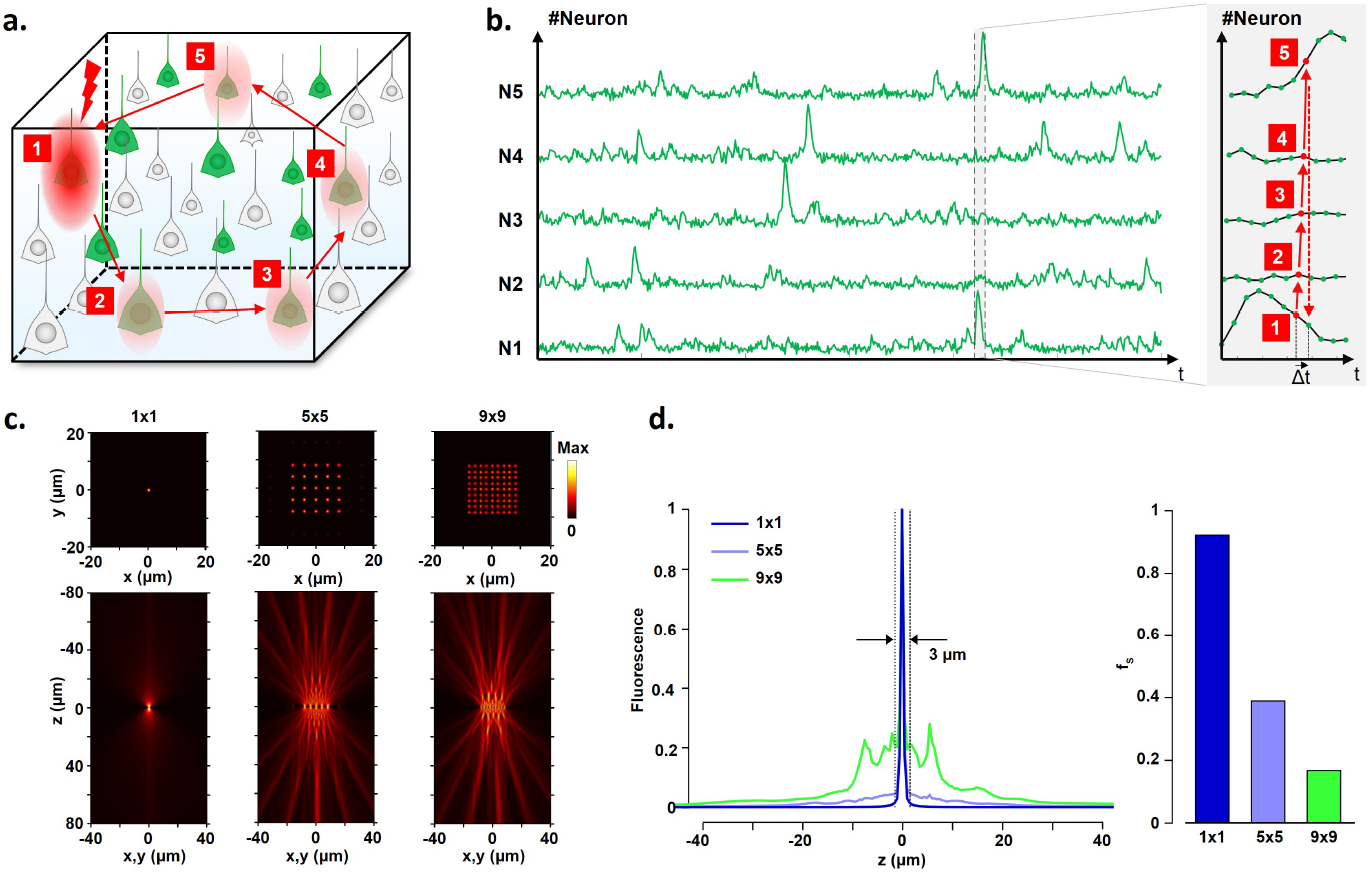
a. Schematic representation of random access 3D recording of a group of neurons labeled with neuronal activity indicator, e.g. for intracellular calcium concentration, annotated in green. b. Simulated functional traces in each neuron with a zoom highlighting the serial recording process. The delay Δ*t* between two samplings of the same trace is defined as *N* / *f* where *N* is the number of neurons and *f* the sampling frequency. c. (xy) and (xz) visualizations of 1 × 1, 5 × 5 and 9 × 9 grids of holographically-multiplexed PSFs used for two-photon fluorescence excitation. d. left: optical sectioning curves associated with the two photon excitation patterns (focusing objective 25x, NA 1.05); right: bar-plot showing the value of *f*_*s*_ (see *Material and Methods*) corresponding to the excitation patterns.

However, a major limitation of this holographic-based random access recording method is the increased background (see Figure 1d). This is primarily due to the larger background arising from the total sample volume, in particular the superficial layers in scattering tissues, for a given signal at the focal plane, as the PSF is holographically enlarged at the focal plane. In addition, holographically-shaped excitation patterns generate interference-induced “hot spots” in the vicinity of the focal plane [13, 14] (see 1). Together, the background generated by these hot spots and by the sample volume decreases the SNR of the recordings and may compromise the specificity of the detected signals. To mitigate the SNR reduction induced by the increased background and to improve signal specificity, activity recordings have been implemented in very sparsely labeled samples [5, 11, 15]. However, this approach inherently limits the possibility of capturing activity across extended neuronal networks of close-by cells. A method to perform random access recordings in densely labeled tissues using holographically-shaped PSF with AODs is thus lacking.

To address this issue, we aim at coupling temporal focusing with AODs [16, 17]. Indeed, temporal focusing (TF) has been successfully demonstrated in association with liquid crystal spatial-light modulators (lc-SLMs) to improve axial resolution by removing Talbot’s periodic intensity patterns around the focal field [13]. However, unlike lc-SLMs, AODs introduce specific spatio-temporal distortions on the excitation beam, such as angular dispersion (AD), group-delay dispersion (GDD) and pulse-front-tilt (PFT) [18], which present unique challenges for integrating TF with AODs. To date, these distortions have prevented the coupling of TF with AODs.

In this study, we present the implementation of TF with AODs to minimize background contributions, supported by both numerical and experimental studies. First, we highlight the consequences of AOD-induced spatio-temporal distortions in the presence of temporal focusing using simulations based on Kostenbauder formalism [19]. Following numerical validation highlighting the possibility to correct the pulse front tilt by an acousto-optic modulator (AOM) added at distance to the AODs, we experimentally demonstrate the recovery of near transform-limited pulses with TF in an AOD-based scheme and significant improvement in optical sectioning in the presence of holography.

## 2. Material and Methods

### 2.1. Kostenbauder formalism

The ray-tracing simulations were performed using the 4 × 4 matrix Kostenbauder formalism to model spatio-temporal distortions - such as PFT, angular dispersion and GDD - experienced by the beam as it propagates through optical elements [19]. Specifically, the basic optical elements and operations used here are represented by the following matrices: ℳ_*lens*_ (*f*) the lens matrix, where *f* is the focal distance of the lens, ℳ_*grating*_ (*λ, β*_*i*_, *β*_*m*_) the grating matrix, where *λ* is the wavelength of the chromatic component, and *β*_*i*_, *β*_*m*_ the angles of the incident and exiting beam, respectively, and ℳ_*propagation*_ (*z, n, k*”) the propagation matrix, where *z* is the propagation distance, *n* the refractive index and *k*” the group velocity dispersion (GVD) of the propagation medium. These matrices can be written as:

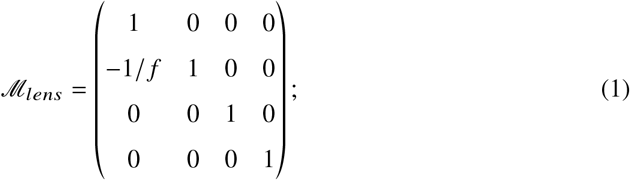

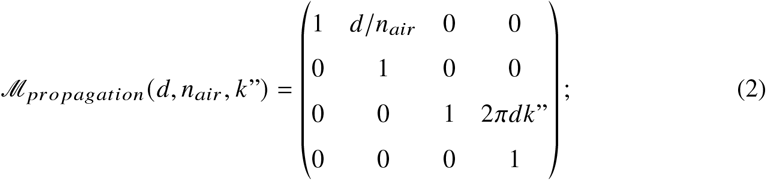

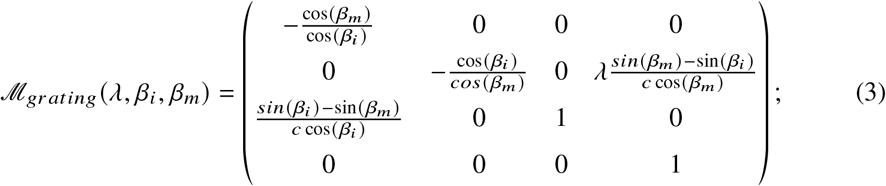

The angle of the incident beam impinging onto the grating is defined as : *β*_*i*_ = −*asin* (*mλ*_0_ /Λ) and the angle at which the beam exits the grating is *β*_*m*_ (*λ*) = *asin* (*mλ* /Λ +sin (*β*_*i*_)), with Λ the spatial constant of the grating. The unpublished AOD matrix, noted ℳ_*AOD*_, was derived following an approach identical to that used to find the grating matrix in [19]. Each matrix element was determined by assessing the variations of the output beam in response to elementary perturbations (translation, rotation, spectral shift, temporal delay) of the incident beam. The AOD and AOM matrices are written as:

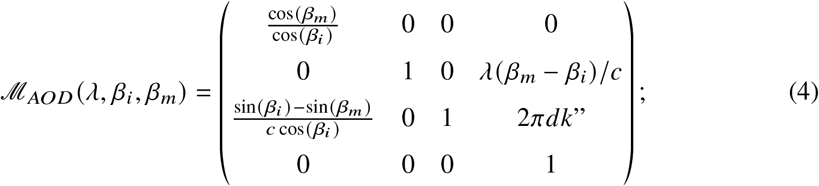

Bragg’s equation is written as follows: 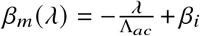 for the AOD and 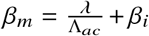 for the AOM (as the AOM is oriented such that the ultrasonic wave propagates in the opposite direction to that in the AOD). These basic elements were used to describe four optical configurations (see Supplementary Figures 1,2): (1) a standard AOD-AOM configuration without TF, as described in [11, 20], which is used e.g. in 3D-CASH for 3D scanning and holographic shaping of the PSF, (2) a standard TF-only configuration without AOD, (3) a TF-AOD configuration without AOM, (4) a TF-AOD-AOM configuration. All configurations include a first telescope for initial beam shaping, a final telescope as an optical relay to the objective back focal plane and propagation through an objective, described as ℳ_*telescope*,1_, ℳ_*telescope*,2_ and ℳ_*objective*_, respectively, and described below:

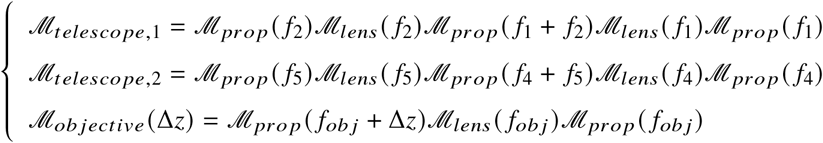

Here, Δ*z* corresponds to the axial distance to the image focal plane of the objective. In the standard AOD-AOM configuration (1), the AOD is positioned at a distance *d* after the AOM, in a plane conjugated to the objective’s back focal plane (BFP). For the TF-configurations (2-4), the grating is placed in a plane conjugated to the objective’s focal plane, while its Fourier plane, in the image focal plane of the lens *ℒ*_3_, is conjugated to the objective’s BFP by the final telescope ℳ_*telescope*,2_. The TF-AOD configuration (3) extends the simple TF-configuration (2) by introducing an AOD in the plane conjugated to the objective’s BFP. In the TF-AOD-AOM configuration (4), an AOM is added to the configuration (3), at a distance *d* before the AOD. Using the matrices previously defined, the four optical configurations were described as the matrix products:

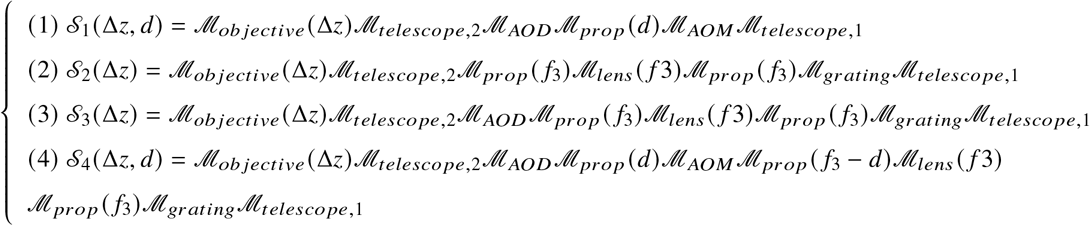

To replicate experimental conditions, the following parameters were implemented in the simulation: the lenses focal lengths *f*_1_ = 75mm, *f*_2_ = 200mm, *f*_3_ = 700mm, *f*_4_ = 300mm, *f*_5_ = 200mm, *f*_*obj*_ = 20mm, the grating with 600-grooves/mm, the AOD and AOM central frequency *f*_0_ = 80MHz, and their frequency bandwidth Δ *f* = 30MHz. We considered different materials: propagation in the air *n*_*air*_ = 1 and *k*”_*air*_ = 0 s^2^/m, TeO2 *n*_*TeO*2_ = 2.27 and *k*”_*TeO*2_ = 5.01 × 10^−25^ s^2^/m. The beam has a diameter 2mm (FWHM), a central wavelength *λ*_0_ = 920nm and a pulse duration *τ*_0_ = 150fs.

### 2.2. Holograms

Holograms were generated using the Gechberg-Saxton algorithm [21], implemented in a phase-only configuration as previously shown in [11]. Four sets of holograms were generated: i) “line with TF”: an horizontal (x-axis) line obtained by focusing the beam along the y-axis and with TF along the x-axis, multiplexed by holography along the y-axis (1 - no multiplexing-, 5 and 9 times); ii) “line without TF” : an horizontal (x-axis) line obtained by focusing the laser along the y-axis only and multiplexed by holography along the y-axis (1 - no multiplexing-, 5 and 9 times); ; iii) “holographic line”: an horizontal (x-axis) line obtained by holography and multiplexed along the y-axis (1 - no multiplexing-, 5, 9 times); iv) “single spot-PSF”: a diffraction-limited PSF multiplexed by holography as a two-dimensional square grids of PSFs (1 × 1 - no multiplexing -, 5 × 5 and 9 × 9). Holograms can be defocused by adding a parabolic profile to the phase, and this parabolic curvature can be added independently on the x or y-axis.

### 2.3. Experimental set-up

The excitation source was a Carbide optical parametric amplifier *(OPA-TW-HP model, Light Conversion)* used at a fixed wavelength of 920nm for all experimental measurements. Power control was achieved using a half-wave plate *(AHW P*10*M* −980, *Thorlabs)* combined with a polarizing beamsplitter cube *(PBS123, Thorlabs)*. The beam was spatially collimated at the laser output with a 1:1 telescope, and pulse duration was measured using an autocorrelator *(IR Carpe, APE)* inserted after the beam collimator. We used a 13mm aperture AOM *(AA*.*MTS/A15@720-920nm, A*&*A, Orsay, France)* and AODs pair *(A*&*A, AA*.*DTS*.*XY/A15@720-920nm)*. The ultrasonic wave frequency and amplitude were controlled in the AOM and in the AOD using a Colibri controller (*Karthala*) and a program written with Labview *(National Instruments)*. At 920nm, the AOD central frequency and bandwidth were respectively 80MHz and 30MHz. The AOM central frequency calculated to be 100MHz, was offset by −12MHz to obtain an optimal angular compensation at the center of the field of view [18].

Two beam shaping settings were implemented for the different optical configurations: in the standard AOM-AOD (no-TF) configuration, the beam was symmetrically expanded to cover the AOD aperture using a spherical magnification telescope *(f*_1_ = 75*mm, AC254-75-B-ML; f*_2_ = 200*mm, AC254-200-B-ML)*; in the “line-TF” configuration”, asymmetric magnification along the y-axis only (vertical axis) was achieved using a cylindrical telescope *(f*_1_ = 75*mm, LJ1703RM-B; f*_2_ = 200*mm, LJ1653RM-B)*. In the standard AOM-AOD (no-TF) configuration, the grating was bypassed using kinematics mirrors mounted on magnetic bases. The AOM, mounted on a 45°-inclined rotation stage attached onto a rail to facilitate translation along the optical axis, was placed at a distance *d* from the X-Y AODs pair. A mirror was placed between the AOM and the AOD to flip horizontally the beam PFT direction. A relay telescope *(f*_4_ = 300*mm, ACT508-300-B-ML; f*_5_ = 200*mm, ACT508-200-B-ML)* conjugated the AOD plane to the BFP of a Leica 10×, NA 0.3 objective. In the “line-TF” configuration, a 600-grooves/mm blazed grating (*GR25-0610*) was placed at the BFP of a cylindrical lens (*f*_3_ = 700*mm, LJ1836L1-B*) which focuses horizontally the spectral components of the beam onto the AOD plane. For the detection scheme, a fluorescently-coated slide was imaged onto a camera *(acA2040-55um, Basler)* using a Nikon 10×, NA 0.45 objective and a tube lens *(f*_6_ = 150*mm, AC254-150-A-ML)*. For assessing optical sectioning profiles, the entire detection assembly was mounted on a rail fixed onto a motorized translation stage to enable axial scanning of the excitation patterns.

### 2.4. Acquisition and analysis of the optical sectioning curves

To assess the axial confinement of each excitation pattern, we performed three-dimensional 2P fluorescence imaging and extracted associated optical sectioning profiles. For this, the detection scheme was translated along the axial direction by Δ*z* = 5*μm* steps to capture the 2P-excitation fluorescence intensity emerging from the excitation of the fluorescent slide for each axial position. A 350 × 350-pixel region was then defined around the 2P fluorescent pattern intensity barycenter determined at the focal plane. Based on this reduced domain, the 3D z-stack was then redefined into a 3D subvolume z-stack. Finally, the 2P fluorescence intensity was plane-averaged and plotted as a function of the axial position (z) to generate the optical sectioning curves S(z).

### 2.5. Background contamination

To estimate the amount of background contamination of the on-focus signal, we defined the “*signal factor*” *fs* from the optical sectioning curve as follows. The signal *F*_0_ is defined as 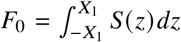, where *X*_1_ is the half width at 1/*e*^2^ of the optical sectioning curve without multiplexing, the background *B* defined as 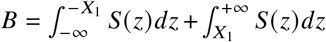 and *fs* as *F*_0_/(*F*_0_ + *B*), such as *fs* varies from 0 (very large background compared to the signal) to 1 (no background).

The presence of the background is detrimental for the recording SNR, as it increases the shot noise, thus making the detection of activity events more challenging.

## 3. Results

### 3.1. Ray-tracing approach based on Kostenbauder formalism

To understand the consequences of spatio-temporal distortions introduced by AODs in the presence of TF and identify a promising TF-configuration with AODs, we performed ray-tracing simulations based on the Kostenbauder matrix formalism [19]. This approach was adopted here for its capability to describe light propagation through spatiotemporally dispersive media [22], specifically to model spatio-temporal distortions including PFT, angular dispersion and GDD. This formalism is an extension of the classical ABCD matrix transformation used in geometrical optics to describe the paraxial propagation of light beams through systems of optical elements in a two-dimensional plane containing the optical axis. In the Kostenbauder formalism, rays are represented as 4 dimensional vectors (*x, θ, t, v*), with *x*, the distance of the ray from the optical axis, *t* the relative delay of the ray to a chosen reference ray, usually the center ray traveling along the optical axis, and *v*, the optical frequency. Optical elements are represented as 4×4 matrices performing linear transformations on these vectors. While Kostenbauder matrices of basic optical elements, such as lenses, mirrors gratings, prisms, etc., were given by Kostenbauder [19], the AOD (and AOM) matrix was derived in this study (see *Material and Methods*) enabling us to test different optical layouts with regard to the effects of spectral dispersion by acousto-optic light diffraction.

Introducing TF into an AOD system raises specific difficulties for the following reasons (Figure 2c). Firstly (Figure 2c top), angular dispersion in AODs following Bragg diffraction will result in significant defocusing in the presence of TF. Secondly, as the combination of AODs with TF spatially separates the different spectral components of the pulse at the AOD input (Figure 2c bottom), the PFT created in the AODs by Bragg diffraction results in GDD in this case. To quantitatively estimate the magnitude of these two effects and analyze how to compensate for them, we modeled the beam propagation in four optical configurations (see Supplementary Figure 1) using the Kostenbauder matrix formalism: (1) AOD-AOM (no TF), (2) TF-only, (3) TF-AOD and (4) TF-AOD-AOM. These simulations provided valuable insights into the spatio-temporal distortions associated with each configuration, in particular those including TF. **AOM-AODs configuration (1)**. This configuration mimics the basic layout of an acousto-optic spatial light modulator (Supplementary Figure 1a) which doesn’t include TF [11, 18, 20, 23]. The role of the AOM in this configuration consists in compensating the spectral angular spread of the transform-limited beam after Bragg diffraction in the AODs, as the diffraction angle is a linear function of the wavelength (law of Bragg diffraction). By setting up the AOM to create angular dispersion of exactly the same magnitude but opposite sign, the AOM-AOD pair collimates the spectral pulse components resulting in an angular dispersion-free output beam (Supplementary Figure 1a), transforming into a point-like focus in the objective focal plane (plane P3 in Supplementary Figures 1a,b). The AOM-AOD configuration also achieves full compensation of Bragg diffraction-induced PFT, as the PFT introduced by Bragg diffraction in the AOD is fully compensated by an opposite PFT in the AOM (Supplementary Figure 1c). However, given the presence of angular dispersion in the space between AOM and AOD, spectral phase delays are introduced through differences in optical path lengths (Supplementary Figure 1a,c). The AOM-AOD pair therefore will produce negative GDD, in analogy with common grating/ prism compressors [24, 25], eventually preventing a full temporal overlap of the spectral components of the pulse in the focal plane (Supplementary Figure 1c, inset). Pulse broadening is expected here because the simulated system lacks normal dispersion by optical materials contributing positive GDD [24, 25]. In real systems, therefore, transform-limited pulse width is restored by tuning the AOM-AOD distance such that the resulting chirp counterbalances the intrinsic system GDD [25], thus functioning as a pre-chirper, yet with the advantage of no additional optics needed.

**Fig. 2.**
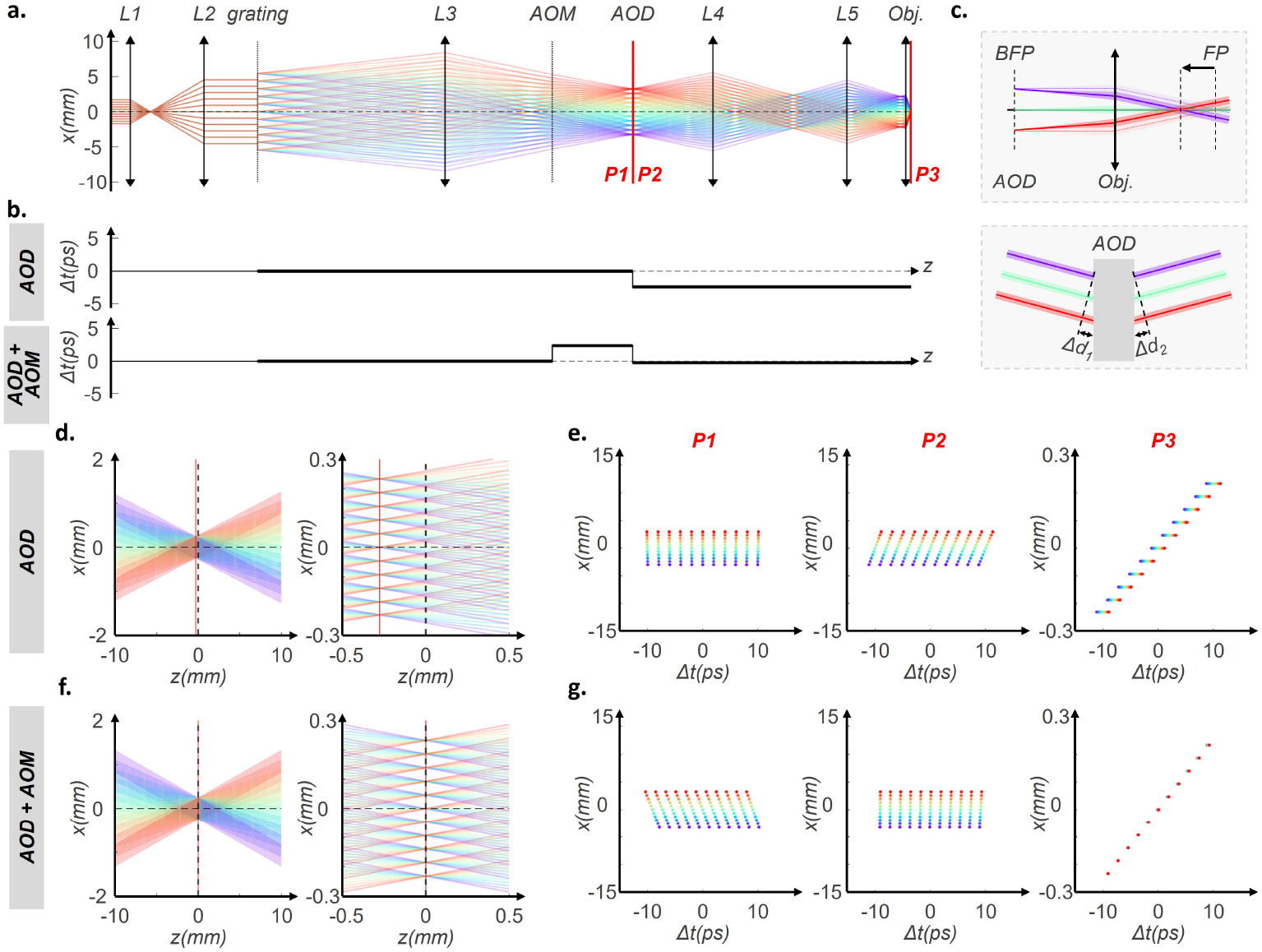
Ray-tracing based on the Kostenbauder approach. a. Ray-tracing in the (xz) plane with x, the TF axis, and z, the optical axis, along the set-up. b. Temporal delay between the spectral components of the beam (1/e^2^). c. Schematics of the defocus induced by angular dispersion (top) and GDD induced by pulse-front-tilt (bottom). d. Left: Ray-tracing in the focal area with the objective approximated as a thin lens of 20 mm focal. Right : Zoom near the image focal plane. e. x-t coordinates plots in plane P1 before AOD, P2 after AOD and P3 at the focal points considered as the position where spectral components cross at the same x position. In the TF-AOD configuration (3), the spatio-temporal distortions result in a defocus and a pulse broadening in duration. In the TF AOD - AOM configuration (4), the spatio-temporal distortions are compensated for, resulting in no defocusing and no GDD at the focal plane (albeit a small residual GDD component uncompensated for by not explicitly accounting for lens material-induced GDD).

**TF-configuration (2)**. The configuration reproduces the line scan version of TF [2, 16], where the dispersive axis of the grating is conjugated with the focal plane of the objective by means of a cylindrical lens *ℒ*_3_ which focuses the spatially dispersed monochromatic beamlets onto a plane conjugated to the objective’s BFP (Supplementary Figure 1d,e). As expected, in the absence of dispersive elements other than the grating, this configuration showed no PFT and no AD-induced GDD, while producing a line focus through a rapid (20 ps) point scan driven by the linear phase shifts along the line introduced by the inclined illumination of the grating (Supplementary Figure 1f). **TF-AOD-AOM configuration (4)**. The configuration (Figure 2b and Supplementary Figure 1j) combines the two above configurations **(1)** and **(2)**. This combination gives rise to two major new phenomena. First, as the grating disperses the frequency components of the beam across the entrance aperture of the AOD (see Supplementary Figure 1g for a simulation in absence of the AOM in the **TF-AOD configuration (3)**), Bragg diffraction-induced AD produces indeed defocus in presence of TF (Figure 2a, upper panel) which, in absence of the AOM (**configuration TF-AOD (3)**), would shift the line focus away from the focal plane (Figure 2d and Supplementary Figure 1h). Secondly, because of the TF frequency dispersion, Bragg diffraction-induced PFT (Figure 2c bottom panel), usually not particularly harmful in a non TF system, will now generate negative GDD of large magnitude, as expected (**TF-AOD configuration (3)**, Figure 2e and Supplementary Figure 1i). While the defocus effect alone could be eliminated by simple means, e.g. by an auxiliary lens or axial displacement of L3, PFT compensation remains imperative in a TF system. As shown above, PFT is fully corrected in the AOM-AOD pair **AOM-AODs configuration (1)**. The same remains true with added TF. In consequence, we expect TF-induced defocus and GDD to be fully compensated in the TF-AOM-AOD configuration (Figure 2f,g and Supplementary Figure 1 k,i), notwithstanding a residual GDD component (Figure 2g) depending upon the AOD-AOM distance as noticed before in absence of TF. In practice, this residual component can be used here also to cancel the positive GDD of the microscope optics, by the fine tuning of the AOM AOD distance.

Overall, spectrally-resolved ray-tracing using Kostenbauder matrices helped us to characterize the spectral phase delays to be expected from optical path differences in systems including dispersive acousto-optic elements and suggest to use the TF AOM - AOD configuration to compensate for these spatio-temporal distorsions.

### 3.2. Retrieving transform-limited pulse duration in acousto-optic TF

We designed an optical setup for testing the different configurations experimentally (Figure 3.a and *Material and Methods*). A blazed diffraction grating with the dispersive axis aligned to the horizontal x-axis is placed in a conjugate plane to the focal plane of an objective of large focal length to allow for precise measurement of the 3D spatio-temporal structure of the focal light pattern. The angularly dispersed frequency components of the laser are focused into the XY-AOD pair by the *L*3 cylindrical lens (x-axis). The AOM is introduced between *L*3 and the AOD and mounted onto an optical rail to adjust the AOM-AOD distance. The AOM is oriented at 45^°^ between the horizontal (x) and the vertical (y) axes, such that the acoustic wave in the AOM is counter-propagatin<u>g</u> with regard to both active axes of the XY-AOD and with its acoustic frequency fixed at around 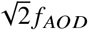, where *f*_*AOD*_ is the central frequency of the AODs bandwidth, to compensate for angular dispersion in both x and y-axis [10, 18]. All measurements were performed at 920nm.

**Fig. 3.**
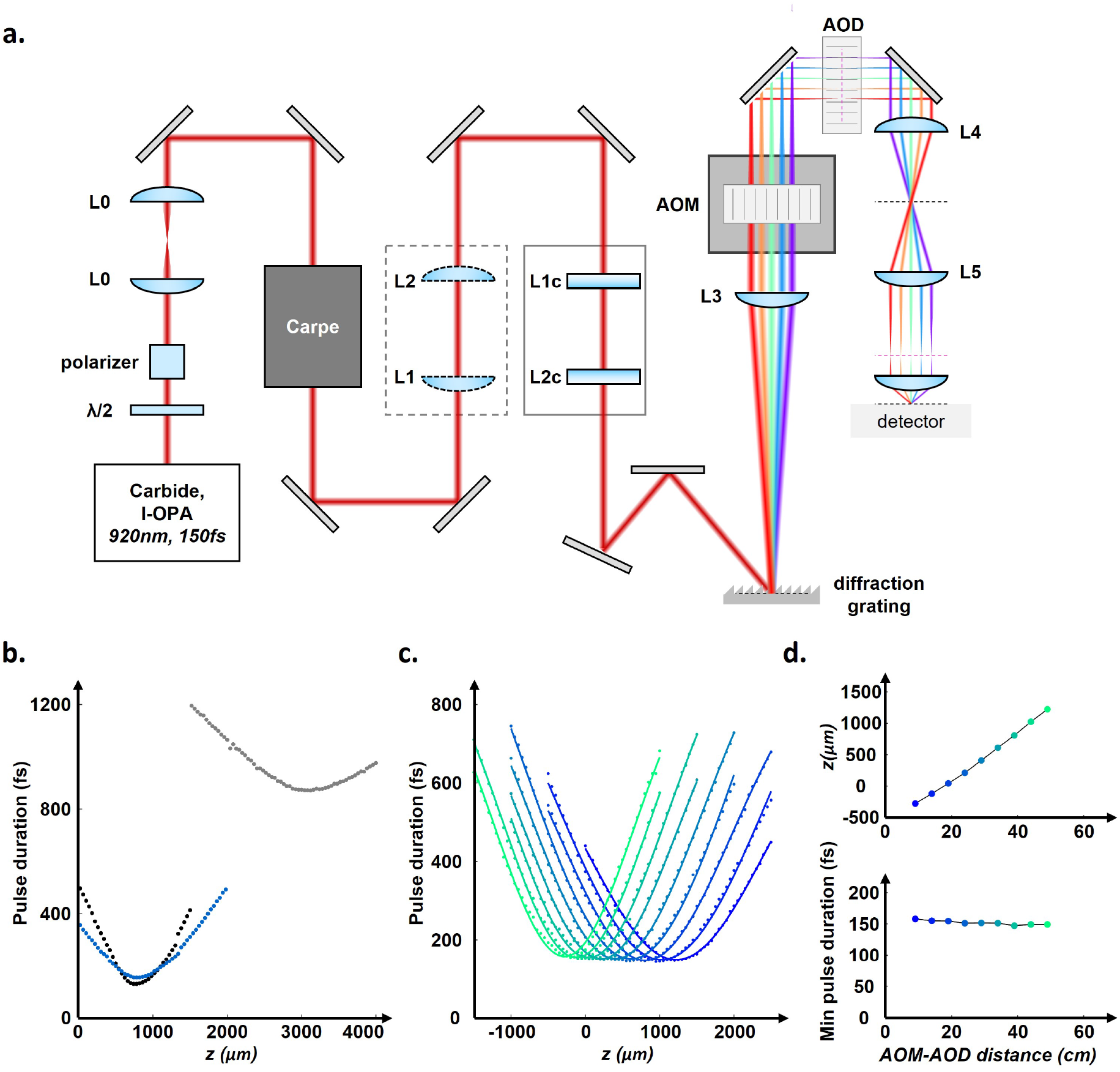
a. Schematic of the experimental set-up detailed in the *Material and Methods* section. b. Pulse duration (FWHM) curves measured in the vicinity of the focal region in presence of TF and in the absence of AOD and AOM (configuration 2, black curve), with AOD only (configuration 3, grey curve) and with both AOD and AOM (configuration 4, green curve). The strong defocus and the large increase in pulse duration observed in the presence of the AOD alone can be corrected by adding the AOM prior to the AOD. c. Curves of pulse duration acquired for different AOM-AOD distances along the optical axis. The axial position where the pulse duration is minimum for a given AOM-AOD distance varies linearly with the axial position of the AOM, and the value of the minimum pulse duration remains constant. All pulse duration measurements were performed with the objective replaced by a 50mm focal length lens.

We first assessed the capability to retrieve transform-limited pulses in the TF configuration with the AOM-AOD pair to confirm the full correction of spatio-temporal distortions at the center of the field of view (FOV). To achieve this, we took systematic measurements of the pulse duration along the optical axis within a range of positions around the temporal focus. The external detector of a Carpe autocorrelator was mounted on a computer-controlled translation stage with nanometer-scale precision to enable axial measurements of pulse duration. Figure 3.b shows the measurement of pulse duration as a function of the axial position of the detector for the three previously described TF configurations: TF only without AOD (configuration (2)), TF with AODs alone (configuration (3)), and TF with both AOM and AODs (configuration (4)). The axial dependence of the pulse duration in the TF only configuration (with AOM and AOD removed) was fitted to determine the minimum pulse duration, which was found to be around 130fs. In contrast, the TF-AOD configuration introduced a significant defocus of 2.3*mm* and increased the minimum pulse duration to 870fs (see Figure 3.b). In accordance with the pulse ray-tracing in the case of TF-AOD (Figure 2.e and Supplementary Figure 1), we assigned the pulse broadening to

Bragg diffraction-induced PFT of the spectrally dispersed beam causing spectral phase delays and thus GDD of considerable magnitude (−6.4 × 10^5^ fs^2^ in the simulation). The observed defocus in TF-AOD, on the other hand, is not revealed by the pulse ray-tracing, as ray-tracing only accounts for spectral delays without explicitly modeling the associated temporal phase. Indeed Durst et al., using a pulse-diffraction model, showed that low to moderate GDD prompts a pure linear defocus with little effect on the pulse width, whereas both nonlinear defocus and pulse broadening dominate at large GDD [26]. The strong increase in pulse duration in TF-AOD suggests that PFT-induced GDD exceeds the regime of linear defocus (Figure 3.b). While the total GDD created by the addition of the AODs must also account for intrinsic GDD of the optical material itself (TeO2), we estimated this contribution to be of the order of 9 × 10^3^ fs^2^ [18] and thus about two orders of magnitude lower in absolute value than PFT-generated GDD. Consequently, we assign the pulse duration increase mostly to PFT-induced GDD. Finally, adding the AOM prior to the AOD restores the transform-limited pulse, while also almost entirely eliminating the defocus (see Figure 3b). This result also provides further evidence that most of the GDD indeed originates from PFT with only minor contribution by normal dispersion of the optical material itself. It appears however that the temporal profile around the minimum distance is slightly broader in TF-AOM-AOD configuration than TF-only and the minimum of pulse duration is marginally higher. This may be due to AOM vignetting of the extreme spectrum components in the dispersed beam. Overall, these results validate the TF-AOM-AOD configuration to achieve full compensation of spatio-temporal distortions in the AOD. Finally, it is important to note that this is effectively due to the positioning of the AOM between *L*3 and the AOD, where the lateral distance between the monochromatic beams is similar. If the AOM were positioned before *L*3, the PFT-induced GDD introduced by the AOD would not be fully corrected by the AOM.

Next, we explored the effect of the AOM-AOD distance on pulse duration. Figure 3.c shows the pulse profiles for a set of AOM positions along the optical axis, and Figure 3.d recapitulates the pulse duration minima values and their associated axial positions for these profiles. Pulse duration is found to remain minimal for all positions, while the defocus varies linearly with the position of the AOM. This result is consistent with the generation of GDD from the angular dispersion after the AOM, which varies linearly with the distance between the AOM and the AOD. As previously described, the angular dispersion introduced by the AOM (regardless of the separation of the chromatic components) is compensated for by the AOD but causes a fixed delay to the spectral components of the beam. This delay depends on the distance between the AOM and the AOD, in a compressor-like manner, and contributes to the global GDD. As this GDD source is low (within the range defined by Durst), it results in a defocus without significant pulse broadening. Again, in practice it can be used to compensate the microscope positive GDD.

### 3.3. AOD-controlled wavefront parabolic curvature ensures complete overlap of the spatial and temporal foci within the FOV

As the TF AOM-AOD configuration (4) enables the recovery of a nearly transform-limited pulse at the FOV center, we next characterized the axial extent of the microscope PSF in the presence of temporal focusing, with two main issues. First, we wondered whether the temporal focus in the grating plane and the spatial focus in the perpendicular plane superpose axially. Secondly, we wondered whether this superposition holds within the field of view (FOV), given that the AOM–AOD configuration only corrects angular dispersion at its centre.

The first step was thus to ensure that the temporal and spatial foci were properly aligned at centre of the FOV, where angular dispersion is fully compensated. In principle, proper superposition of the two foci can be achieved in two ways: either by displacing the grating along the optical axis to axially shift the temporal focus, or by introducing curvature in only the Y-AOD to axially shift the spatial focus accurately (see *Material and Methods*). Figure 4.a shows (xy) and (xz) views of the 2P fluorescence stack acquired by translating a thin fluorescent slide through a line pattern obtained by temporal focusing. Adding a small 5 *μm*-defocus reveals a thin line in the temporal focus plane and a single axial focus in the (xz) projection, thus optimizing the overlap between the temporal and spatial foci, by correction of a residual defocus. In contrast, introducing a 30 *μm* defocus shifts the spatial focus away from the temporal focus. In this case, the fluorescent line appears thicker in the temporal focus plane and a double focus becomes visible in the (xz) projection. To confirm this observation, we applied a series of curvatures to the Y-AOD and acquired the corresponding 2P fluorescence stacks to explore the capability of shifting the spatial focus over a larger axial range. Figure 4.b shows the normalized optical sectioning profiles resulting from averaging the (xy) 2P fluorescence for this series of spatial defocus. While the position of the fluorescence peak associated with the temporal focus remains unchanged, the position of the fluorescence peak originating from the spatial focus can be shifted along the optical axis by applying curvature to the Y-AOD. Both peaks coincide precisely when a defocus of 5 *μm* is applied, in agreement with the previous result shown in Figure 5a. This shows that both foci can be perfectly superimposed by taking advantage of the possibility to add an accurate amount of parabolic curvature with the Y-AOD.

**Fig. 4.**
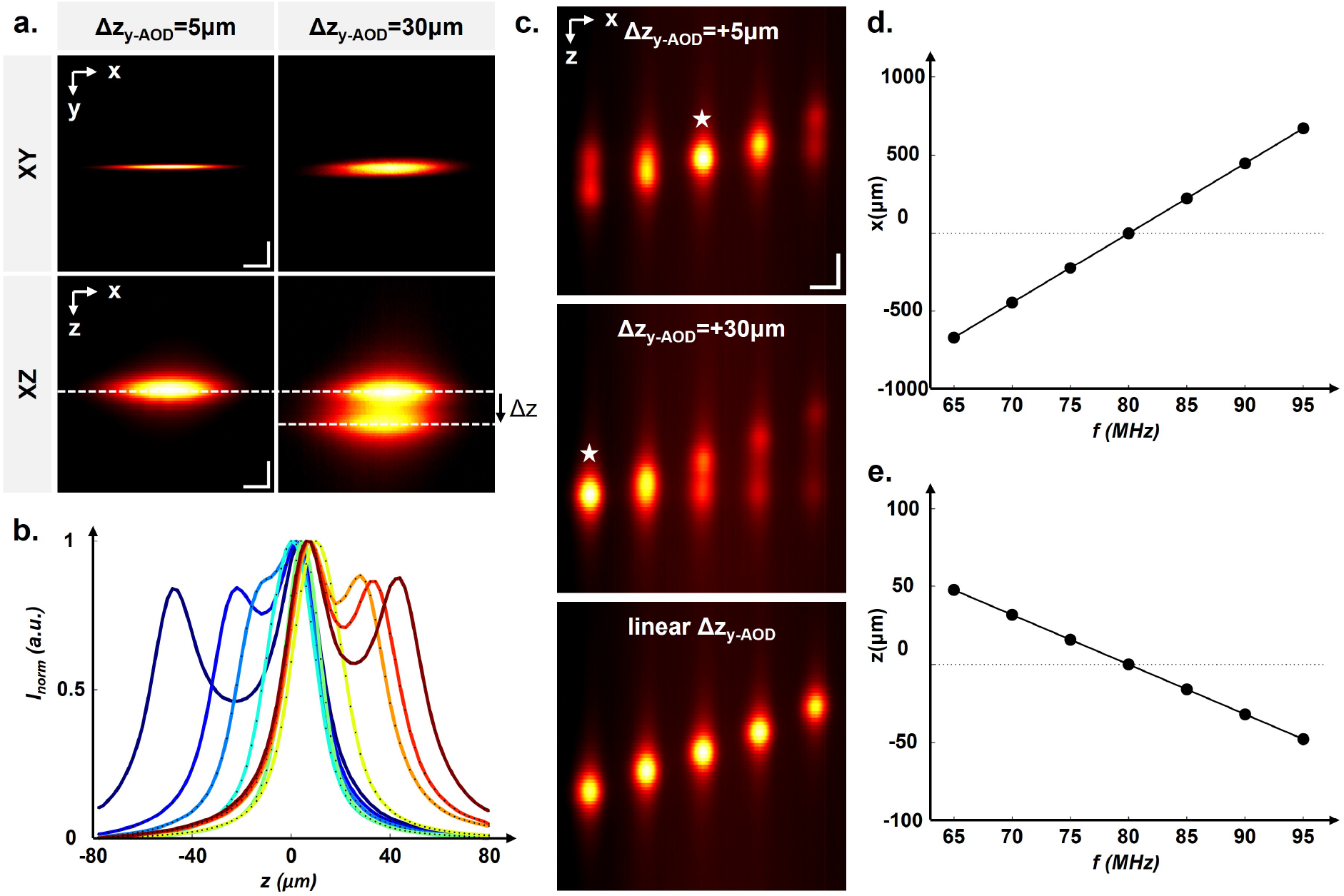
a. (xy) and (xz) 2P fluorescence images from a fluorescent slide with a 5 *μm* (1st column) and 30 *μm* (2nd column) parabolic curvature added to the Y-AOD. The first column shows the full overlap of the spatial and temporal foci, while the second column shows a shifted spatial focus with respect to the temporal focus. Scales: 20*μm* in x-, y- and z-directions. b. Optical sectioning curves obtained by averaging (xy) intensity from z-stack obtained for a series of parabolic curvature applied on the Y-AOD. c. (xz) fluorescent images with 5 patterns spared along the x-axis in the FOV for a parabolic curvature applied to Y-AOD of 5 *μm* (upper panel), 30 *μm* (middle panel) and which increases linearly with the x-position (lower panel). While a fixed parabolic curvature offset on the Y-AOD only corrects the shift between the temporal and spatial foci at a specific x-position, the application of parabolic curvature that varies linearly across the x-axis fully compensates the shift between the foci along the whole x-axis. Scales: 100*μm* in x- and 20*μm* in z-direction. d. and e. Dependence of the lateral (d) and the axial (e) positions of the temporal focus along the x-axis obtained using Kostenbauder simulations. These results confirm the presence of an axial shift of the temporal focus as a function of the x-position originating from residual uncompensated angular dispersion in the FOV.

**Fig. 5.**
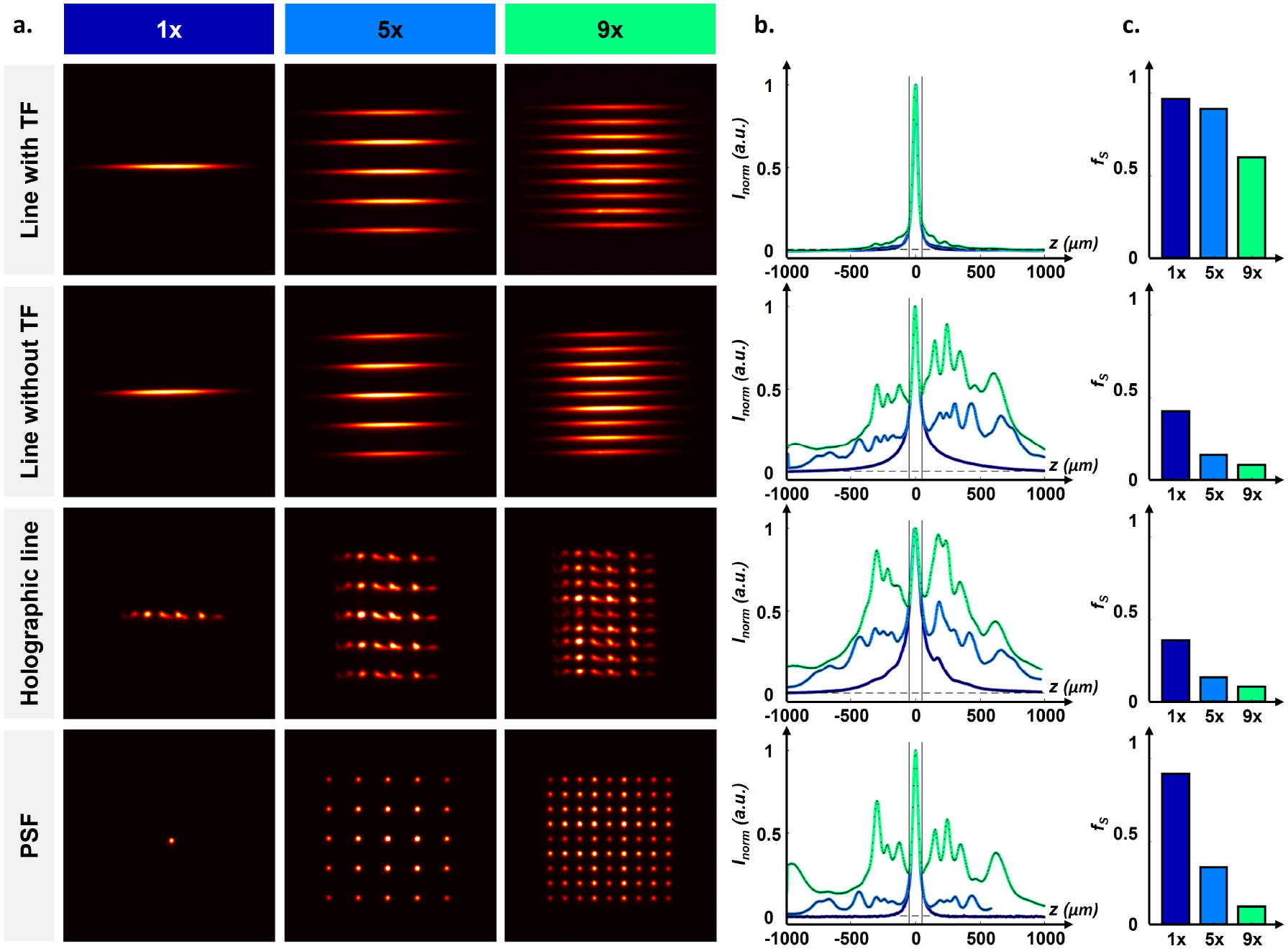
a. Images of two-photon fluorescence of the excitation patterns in the focal plane. The first column shows the original pattern; the second column shows the pattern multiplexed 5× along the y-axis; and the third column shows the pattern multiplexed 9× along the y-axis. For the last row the pattern is multiplexed along the x and y direction. b. Optical sectioning curves obtained by averaging the x-y fluorescence collected from the patterns displayed in (a). c. Bar plot showing the signal fraction *f*_*S*_ calculated from the optical sectioning curves obtained for each pattern. The first line corresponds to a line with temporal focusing along the x-axis; the second line refers to a line without TF; the third line corresponds to a holographic line and the fourth line shows a PSF.

However, this alignment might not hold in the field of view, as the angular dispersion is only fully compensated at its center, which could cause a shift of the temporal focus in the FOV. We explored thus the capability to overlap the spatial and temporal foci within the FOV. Figure 4.c shows (xz) projections for different positions of the TF-line pattern along the x-axis within the FOV. As expected, the spatial focus remains confined to the same plane and the temporal focus undergoes a linear defocus along the x-axis (Figure 4.c upper panel). Adding a 30 µm defocus to the Y-AOD axially shifted the plane of spatial focusing, enabling both foci to overlap on the left of the FOV (Figure 4.c middle panel). Finally, the perfect overlap of both spatial and temporal foci across the x-axis could be obtained by applying a tunable curvature to the Y-AOD (Figure 4.c. lower panel). It should be noted that this correction can be performed at any point on the x-axis for each successive laser pulse, thanks to the fact that AODs can apply the necessary wavefront both to reach a certain position in the field of view and to correct this deviation for each laser pulse, at the laser repetition rate (40kHz), which is a unique feature of this system.

To understand more quantitatively the axial shift of the temporal focus that increases linearly along the x-axis, we aimed to model it numerically using Kostenbauder simulations (see also Supplementary Figure 2). Figure 4.d shows the lateral and axial positions of the temporal focus as a function of the X-AOD frequency, *f*_*AOD*_, which corresponds to different lateral x-axis positions. They predict an axial defocus of the temporal focus along the x-axis, of typically 7.1*μm* for a 100*μm* lateral shift along the x-axis. This result is consistent with our experimental observations and confirms that this effect results from the angular dispersion, and can be fully compensated only at the FOV center. Residual angular dispersion in the FOV increases linearly with the x position and converts into defocus. This effect is identical to the conversion of angular dispersion into defocus at the FOV center in the presence of an AOD alone.

### 3.4. Applying TF to multiplexed patterns minimizes background contamination

We examined here the effect of the temporal focusing on the axial confinement of excitation patterns generated using AOD-based holography. Specifically, we compared the confinement of the following four patterns: (i) a “line with TF” which corresponds to temporal and spatial foci along the x- and y-axes respectively; (ii) a simple “line without TF” resulting from a spatial focus along the y-axis only but with neither spatial focusing nor TF along the x-axis; (iii) a “holographic line” obtained using a holographic pattern achieved with the X-AOD, a spatial focusing on the y-axis and no temporal focusing; and (iv) a standard “single spot-PSF” resulting from spatial foci along the x- and y-axes (Figure 5.a, first column). All patterns were then multiplexed by holography using the Y-AOD. Patterns were multiplexed either 5 times (Figure 5.a, second column) or 9 times (Figure 5.a, third column), such that all resulting patterns cover a spatial domain of similar lateral size. Additionally, the “single spot-PSF” of the pattern (iv) was also multiplexed along the x-axis using the Y-AOD to extend the pattern laterally along the x direction too (Figure 5.a, last row). These extended patterns are analogous to the ones covering the neuron soma and used in 3D-CASH to record their activity with GECIs [11].

Figure 5.a shows focal plane images of all these 1×, 5× and 9× multiplexed patterns obtained from two-photon fluorescence z-stack acquisitions. In the first two rows, the lines obtained with and without TF exhibit the same homogeneity in the focal plane. In contrast, the line generated by holography in configuration (iii) shows a speckled heterogeneous intensity distribution along the x-axis. The multiplexed PSF grids from configuration (iv) display a regular PSF distribution, albeit with heterogeneous intensity due to the use of phase holography rather than mixed phase and intensity holography [12].

Figure 5.b shows the optical sectioning curves obtained by averaging the 2P fluorescence intensity across the (xy) planes of the 2P z-stack collected from each pattern. To quantify the axial confinement of the excitation pattern, we derived the signal fraction parameter *f*_*S*_ from these optical sectioning curves, as described in the *Material and Methods* section. Figure 5.c shows bar plots of the signal factor *f*_*S*_ for all the patterns.

In the absence of multiplexing (see Figure 5.b), all optical sectioning profiles exhibit PSF-like profiles. As expected, the “line with TF” and the “single spot-PSF” configurations, which present two dimensions of focalization, exhibit improved axial confinement, with *f*_*S*_ > 0.9 (see Figure 5.c). In contrast, the “line without TF” and the “holographic line” configurations present a degraded axial confinement with *f*_*S*_ ≃0.35 − 0.45, due to the presence of only one focalization dimension.

In the presence of multiplexing (see Figure 5.b), 2P fluorescence intensity peaks appear off-focus on the optical sectioning curves in configurations (ii-iv). These intensity peaks result from “hot spots” emerging axially away from the focus, and lead to a drastic decrease in the *f*_*S*_ parameter. Furthermore, increasing the multiplexing density yields increased 2P fluorescence intensity away from the focus. This is due to a larger concentration of hot spots in the vicinity of the focus. In contrast, the profiles from the multiplexed TF-line pattern remain relatively unchanged for a 5× multiplexed pattern, and show a subtle degradation for the 9× multiplexed pattern. Consequently, the *f*_*S*_ factor remains largely preserved for denser multiplexed patterns.

## 4. Discussion

In this paper, we presented the implementation of temporal focusing in an AOD scanning system for two-photon microscopy, in a configuration named 3D-CASH that allows 3D scanning at 40 kHz rate with an holographically-shaped PSF. While 3D-CASH provides uniquely the possibility to record neuronal activity in 3D with millisecond resolution, the increased background contamination of the on-focus signal limits its use to sparsely labeled samples and small neuronal networks.

To implement temporal focusing in this system for reducing the background contamination, we first identify a suitable configuration using a simulation based on Kostenbauder matrices. To model the consequences of the spatio-temporal distortions introduced by AODs, we first derived the Kostenbauder matrix describing the AOD, following the method used for a grating in [19]. We then demonstrated that the introduction of an AOD at the back focal plane of the microscope in presence of temporal focusing has two major consequences, namely a defocus originating from the angular dispersion linked to Bragg diffraction, and a GDD caused by pulse-front-tilt. These two effects both arise from the lateral separation of chromatic components at the AOD plane. We finally showed that adding an AOM prior to the AOD corrects both effects at the FOV center by introducing inverted angular dispersion and PFT. Beyond the application presented here in the context of AOD-based microscopy [11, 18], the Kostenbauder approach could be of interest for characterizing spatio-temporal distortions in other acousto-optic systems, but also when incorporating angular dispersive devices such as liquid crystal spatial-light modulators (lc-SLM).

Experimental measurements show the retrieval of a nearly transform-limited pulse following the introduction of the AOM, thereby validating the correction of both defocus and GDD. Taking advantage of the possibility to introduce controlled wavefront modulation independently in each AOD, we also demonstrated the capability to precisely overlap the temporal and spatial foci at the FOV center. Furthermore, we have demonstrated experimentally and numerically the presence of an intrinsic axial shift of the temporal focus that increases linearly with the distance in the direction of the temporal focusing and that originates from the residual uncompensated angular dispersion in the field of view. Introducing a controlled parabolic curvature to the spatial focus with the corresponding AOD enabled the spatial and temporal foci to overlap accurately at each point in the field of view. The possibility to correct this shift between the two foci at all positions is a striking property of the AODs, as they allow synchronously and at 40kHz to scan the beam in 3D and shape the wavefront.

The introduction of temporal focusing in the AOD system reveals to provide a very significant gain in background rejection, as expected. We showed indeed that an improved background rejection by a factor of 6 can be achieved using an excitation pattern consisting of a line obtained by temporal focusing multiplexed 9 times in the perpendicular direction by holography, as compared to a pattern of similar dimension consisting of 2D array of single spot-PSF obtained by 2D holography. This result opens promising perspective for in vivo neuronal activity recordings, with the possibility to improve significantly the recordings SNR, and thus the ability to detect spikes with high fidelity, and to extend this methodology to densely labeled samples and larger neuron networks, a strong limitation at this time.

To implement this new temporal focusing AOD system for in vivo application, it will require to adapt it to high numerical aperture objectives and to design patterns adapt to the GEVIs and GECIs staining. For that reason, we found that combining temporal focusing along one axis and holography along the other axis may provide more flexibility for the design of patterns as compared to a configuration with temporal focusing in 2D dimensions. The latter configuration is bound to generated disk patterns, which are most probably not optimal when used e.g. with GEVIs indicators labeling the plasma membrane only. Our configuration may facilitate the design of patterns that optimize photons flux, depending on the type of indicators, and recording stability.

## 5. Conclusion

Background contamination is a major limitation of random-access recordings techniques based on a holographically-shaped Point Spread Function. However, combining acousto-optic deflectors, the most powerful device for these approaches, with temporal focusing, a proven method to reject background in digital holography, is not straightforward. Here we have described the complex spatio-temporal distortions faced when coupling temporal focusing with acousto-optic deflectors. We showed that the introduction of an acousto-optic modulator reveals to fully compensate for these complex distortions at the center of the field of view, as in the classical configuration without temporal focusing, where the AOM compensates for angular dispersion due to Bragg diffraction. In addition, taking advantage of the ability of AODs to shape the PSF synchronously with the 3D scans, we showed that the shift between the temporal and spatial foci in the field of view resulting from uncompensated angular dispersion can also be corrected with high accuracy. Coupling temporal focusing in one direction and holography in the perpendicular one, allowed us finally to generate extended two-photon excitation patterns with significantly reduced background excitation, as compared to pure 2D holographic patterns, offering particular benefit to in-vivo random-access recording of neuronal activity.

## Ackowledgments

We would like to thank Vincent Villette, Stéphane Dieudonné, Cathie Ventalon, Sébastien Wolf and the IBENS Imaging Platform for their ongoing support of this work. We would also like to thank Alberto Lombardini, Ruth Sims and Eirini Papagiakoumou for their valuable contributions to discussions about temporal focusing.

## Data Availability Statement

Data underlying the results presented in this paper are not publicly available at this time but may be obtained from the authors upon reasonable request.

