## Supplementary Material for "Temporal Focusing for Enhanced Background Rejection in AOD-Based Two-Photon Serial Holography"

### 1. Ray-tracing on the optical axis based on Kostenbauder simulations

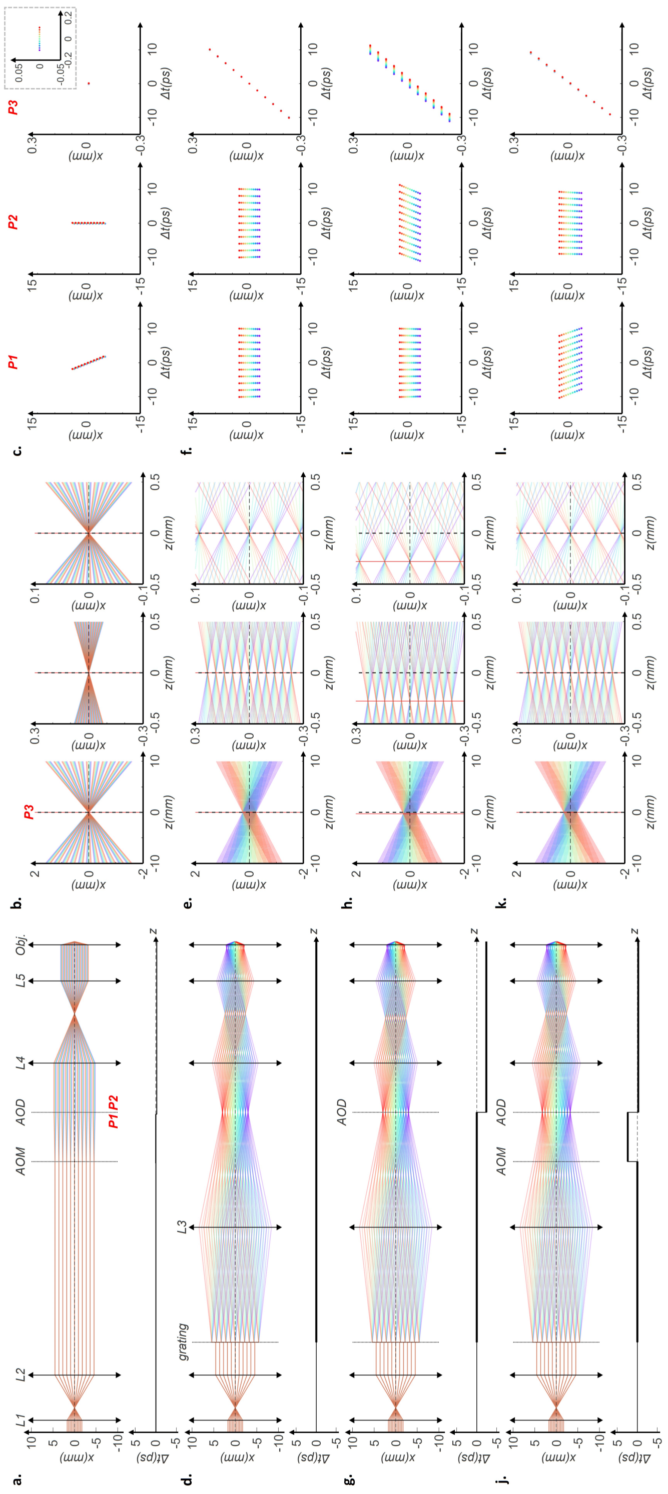


Fig. 1. Ray-tracing on the optical axis in the following configurations: (a) AOM-AOD without TF (configuration 1), (b) TF only (configuration 2), (c) TF-AOD (configuration 3) and (d) TF-AOD-AOM (configuration 4). The first column shows ray-tracing in the (xz) plane with x representing the TF axis, and z the optical axis (shown above). The temporal delay between the spectral components of the beam ($1/e^2$) is shown below. Columns 2-4 show ray-tracing in the focal area. Columns 5-7 represent x-t coordinate plots in planes P1 and P2, before and after the AOD plane, respectively and in P3 at the focal points considered as the position where spectral components cross at the same x position.

### 2. Ray-tracing out-of-the optical axis based on Kostenbauder simulations

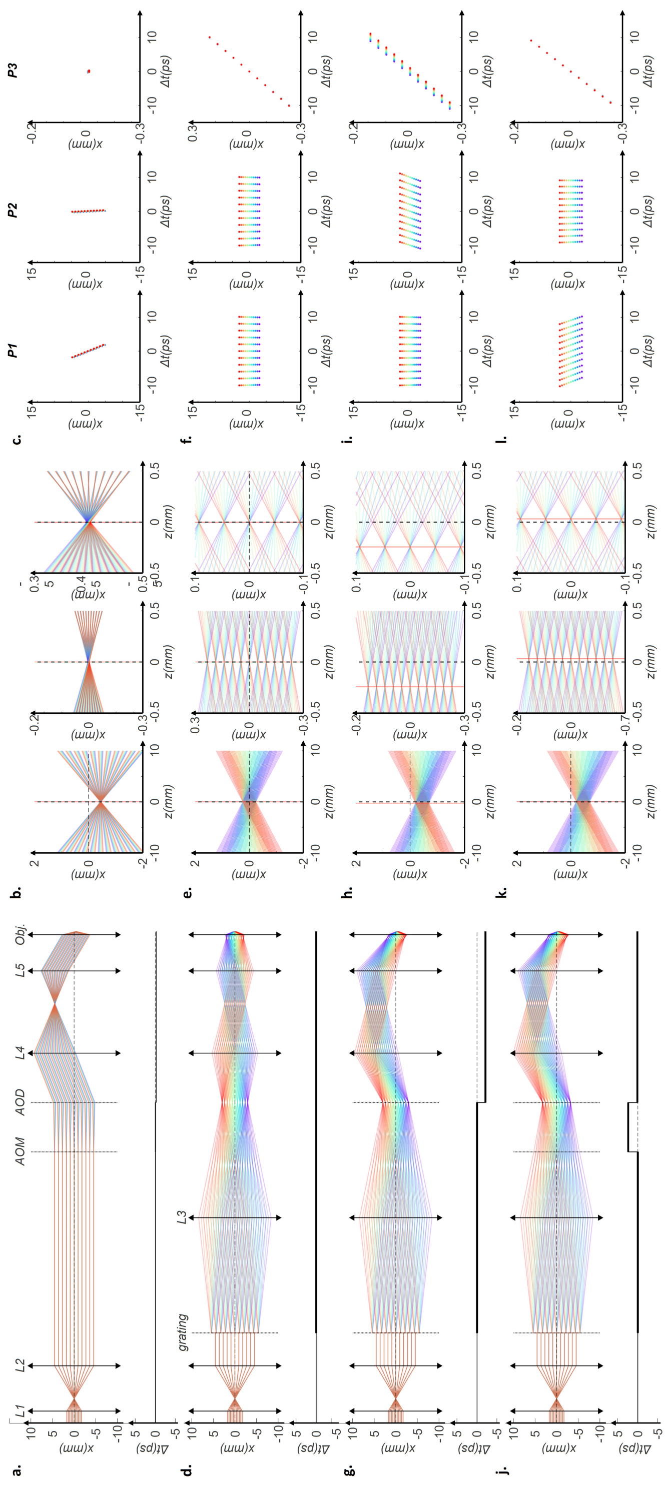


Fig. 2. Ray-tracing out-of-the optical axis in the following configuration: (a) AOM-AOD without TF (configuration 1), (b) TF only (configuration 2), (c) TF-AOD (configuration 3) and (d) TF-AOD-AOM (configuration 4). The first column shows ray-tracing in the (xz) plane with x representing the TF axis, and z the optical axis (shown above). The temporal delay between the spectral components of the beam ($1/e^2$) is shown below. Columns 2-4 show ray-tracing in the focal area. Columns 5-7 represent x-t coordinate plots in planes P1 and P2, before and after the AOD plane, respectively and in P3 at the focal points considered as the position where spectral components cross at the same x position.
